# Cell cycle-dependent chromatin dynamics at replication origins

**DOI:** 10.1101/2021.11.15.468728

**Authors:** Yulong Li, Alexander J. Hartemink, David M. MacAlpine

## Abstract

Origins of DNA replication are specified by the ordered recruitment of replication factors in a cell cycle dependent manner. The assembly of the pre-replicative complex in G1 and the pre-initiation complex prior to activation in S-phase are well characterized; however, the interplay between the assembly of these complexes and the local chromatin environment is less well understood. To investigate the dynamic changes in chromatin organization at and surrounding replication origins, we used micrococcal nuclease (MNase) to generate genome-wide chromatin occupancy profiles of nucleosomes, transcription factors and replication proteins through consecutive cell cycles in *Saccharomyces cerevisiae*. During each G1 phase of two consecutive cell cycles, we observed the downstream repositioning of the origin-proximal +1 nucleosome and an increase in protected DNA fragments spanning the ARS consensus sequence (ACS) indicative of pre-RC assembly. We also found that the strongest correlation between the chromatin occupancy at the ACS and origin efficiency occurred in early S-phase consistent with the rate limiting formation of the Cdc45-Mcm2-7-GINS (CMG) complex being a determinant of origin activity. Finally, we observed nucleosome disruption and disorganization emanating from replication origins and traveling with the elongating replication forks across the genome in S-phase, likely reflecting the disassembly and assembly of chromatin ahead of and behind the replication fork, respectively. These results provide insights into cell cycle-regulated chromatin dynamics and how they relate to the regulation of origin activity.

## INTRODUCTION

Duplication of a cell’s genetic information occurs every cell cycle in S-phase. While DNA replication is restricted to S-phase, the DNA replication program is established earlier in the cell cycle with the licensing of DNA replication origins in G1 [1]. These licensed origins are then activated with an inherent efficiency during S-phase [2]. While many studies have carefully examined the kinetics of DNA replication progression through S-phase [3–7], few have examined the chromatin dynamics of replication origins as cells progress through multiple cell cycles. Instead, most chromatin-based studies have interrogated static snapshots of chromatin from discrete cell cycle phases (e.g. G1) or from an asynchronous population of cells [8–11]. Understanding how chromatin structure and organization change at DNA replication origins as they progress through multiple unperturbed cell cycles will provide important insights into the chromatin features that modulate origin usage and efficiency.

The selection and activation of replication origins involve the recruitment of a series of replication factors in an ordered manner through multiple phases of the cell cycle. In *Saccharomyces cerevisiae*, potential origins, which are defined by an autonomous replicating sequence (ARS) and contain a conserved T-rich ARS consensus sequence (ACS), are recognized and bound by the hetero-hexameric origin recognition complex (ORC) [12,13]. In G1, the Mcm2-7 replicative helicase is loaded in a Cdc6 and Cdt1-dependent manner to form the pre-replicative complex (pre-RC) and license origins for activation [14]. Additional initiation factors, including Cdc45 and GINS, are recruited in S-phase to assemble the active helicase known as the Cdc45-Mcm2-7-GINS (CMG) complex that commits origins for activation [15]. The concentrations of Cdc45 and components of GINS are significantly lower than those of the pre-RC and serve as a rate limiting step in the activation of individual origins [16,17].

Only a small subset of ACS motif matches in the yeast genome are *bona fide* ORC-binding sites, and chromatin architecture is believed to be an important factor in defining origins [8,18,19]. Origins of replication in the budding yeast have a stereotypical chromatin architecture with well positioned nucleosomes surrounding a nucleosome free region (NFR) containing the ACS [8,20]. The chromatin architecture with well positioned nucleosomes appears to be a conserved feature of eukaryotic origins [21–23]. In addition, the inherent initiation efficiency [2] and the time of activation [3,24] of each origin are also thought to be regulated, in part, by the local chromatin environment and the levels of chromatin associated ORC [20,25] and Mcm2-7 [26,27].

The selection and activation of DNA replication origins are tightly coupled with the cell cycle. Concomitant with the recruitment of factors to the origin in G1 to assemble and load the helicase, the NFR expands in a Cdc6-dependent manner to accommodate pre-RC assembly [20,28]. Further, as cells enter S-phase in the absence of primase activity, the disorder or entropy of the origin-flanking chromatin increases markedly, presumably due to helicase activation and nucleosome eviction [29]. Finally, the stability and occupancy of ORC on the DNA throughout the cell cycle are origin dependent and predictive of origin efficiency [25]. The majority of origins exhibit a protected footprint representing ORC and/or the pre-RC in the NFR in both G1 and G2 phases of the cell cycle; however, select origins only exhibit a defined footprint in G1 suggesting a more dynamic or transient interaction with ORC which may be stabilized by pre-RC assembly [20]. The stability of ORC and/or pre-RC components on the DNA is predictive of origin efficiency, with more efficient origins having a protected footprint in both G1 and G2.

To better understand the cell cycle-regulated chromatin dynamics with high spatiotemporal resolution, we generated genome-wide chromatin occupancy profiles [20,30] of chromatin sampled at multiple points throughout two consecutive cell cycles. This approach provides factor-agnostic occupancy profiles of DNA-binding proteins, including nucleosomes, replication factors and transcription factors at nucleotide resolution. We comprehensively profiled the dynamics of protected fragments at the ACS and the organization of ACS-flanking nucleosomes throughout multiple cell cycles. Our study describes the chromatin architecture at individual origins in a synchronized cell population and associates cell cycle dependent chromatin features with origin efficiency, thus providing mechanistic insight into the dynamic interplay between chromatin architecture and origin function.

## MATERIALS AND METHODS

### Yeast culture and cell cycle time courses

W303 yeast strain was used in this study with the genotype *MATa*, *leu2-3*, *112*, *BAR1::TRP*, *can1-100*, *URA3::BrdU-Inc*, *ade2-1*, *his3-11*, *15*. Yeast cells were grown in YPD (1% yeast extract, 2% peptone, 2% dextrose) at 30°C to an OD_600_ of ~0.3 and arrested in G1 phase with α-factor (GenWay) at a final concentration of 50 ng/mL for 2 h. Samples were taken right before release as the “α-factor” time point. Cells were then washed twice in sterile water, resuspended in fresh YPD medium, and samples were collected every 10 min until 150 min post release. For each time point, 40 mL of culture was crosslinked with a final concentration of 1% formaldehyde at room temperature for 30 min, quenched with 0.125 M glycine for 5 min, washed and flash frozen. In parallel, 1 mL of culture was resuspended in 70% ethanol and fixed overnight at 4°C for flow cytometry. Independent biological duplicates were performed.

### Chromatin digestion with MNase and sequencing library preparation

MNase digestion of chromatin and sequencing library preparation were performed as previously described [20,30] with the following modifications: 2 μg of digested DNA was used as input; NEBNext multiplex oligos for Illumina Kit (New England Biolabs) was used in adapter ligation and PCR amplification steps; and PCR reactions were performed with 12 cycles and libraries were purified using Agencourt AMPure XP beads (Beckman). Libraries were sequenced on NextSeq 500 High-Output 25bp PE platform (Illumina).

### Flow cytometry

Fixed yeast cells were washed with water, briefly sonicated, and incubated in 50 mM sodium citrate (pH 7.4) with 0.3 mg/mL RNase A for 2 h at 50°C. Then, 0.6 mg/mL Proteinase K (Worthington) was added and incubated for an additional 2 h at 50°C. Finally, cell pellets were resuspended in 50 mM sodium citrate with 1:5,000 SYTOX green (Invitrogen) and incubated for 1 h at room temperature. Flow cytometry was performed on a BD FACSCanto analyzer, and 30,000 cells were recorded for each sample.

### Sequencing data processing and analysis

All reads were aligned to the sacCer3/R64 version of the S. *cerevisiae* genome using Bowtie 0.12.7 [31]. MNase-seq reads were mapped in paired-end mode with the following Bowtie parameters: -n 2 -l 20 --phred33-quals -m 1 --best --strata -y. Data analysis was performed in R version 3.2.0. All genomic data are publicly available at the NCBI GEO repository with the accession number GSE168699.

Because the position of each MNase-seq fragment could be determined by the coordinate of one end and the fragment length, only reads mapped on the forward strand were kept. MNase-seq data from the biological duplicates were randomly subsampled and merged to reduce bias from MNase digestion, library preparation, and sequencing depth before downstream analysis. For each replicate over all time points, the fewest number of reads for each fragment size (from 20 bp to 250 bp) was identified and used as the subsampling depth. The MNase-seq data for each time point was then subsampled to the above depth per fragment size to assign equal number of reads for each fragment length among all time points. Reads mapped to mitochondrial DNA (chrM) or ribosomal DNA regions (chrXII: 451,575 – 489,469) were excluded. After subsampling, the total number of reads for each time point was ~17 million and ~21 million for Replicates 1 and 2, respectively. The matched time points between duplicates were merged for downstream analysis.

### Quantification of nucleosome occupancy

For each time point, a pileup matrix of fragment size by fragment midpoint position was calculated for the aggregate MNase-seq signal of 8,632 unique nucleosome positions on Chr IV which were mapped by a sensitive chemical mapping method [32]. This matrix represents the approximate size and coverage distribution of MNase-seq reads centered at a canonical well-positioned nucleosome. A two-dimensional kernel was then derived using a bivariate Gaussian distribution parameterized by the marginal means and variances of the matrix [33]. The variance of fragment size dimension (y axis) was set to 1/16 of the original marginal variance and the variance of midpoint position dimension (x axis) was set to 1/4 of the original marginal variance. To quantify the occupancy signal of a nucleosome at a given chromosomal location, a cross-correlation score was computed between the local MNase-seq signal matrix and the model nucleosome kernel.

To correct for replication-dependent DNA copy number variation throughout the cell cycle for a given chromosomal location, the RPKM of all MNase-seq reads for a 1,001-bp window centered at the given position was calculated for each time point, and the ratio over the RPKM of the α-factor time point (G1) was considered as the copy number. For each time point, the nucleosome score for any chromosomal location was normalized by its copy number.

### Quantification of small fragment occupancy

For each chromosome, the midpoint density of fragments smaller than 120 bp was estimated using a Gaussian kernel at a bandwidth of 50. The small fragment occupancy of a given chromosomal location was calculated as the product of density and chromosome length and normalized by the copy number of the given position. To adjust for variations of MNase digestion among samples, the average signal of the aggregate small fragment occupancy within +/− 100 bp around 151 Abf1p binding sites downloaded from http://fraenkel-nsf.csbi.mit.edu/improved_map/p001_c3.gff [34] was calculated for each time point and the reciprocal of which was used as a scale factor.

### Quantification of nucleosome disorganization by Shannon entropy

For a region *X* of size *n* bp, the probability of nucleosome positioning at location *i* was defined as

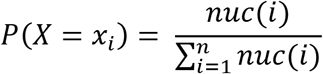

where *nuc*(*i*) is the nucleosome score at position *i*.

The disorganization of nucleosome positioning for region *X* was measured using Shannon entropy

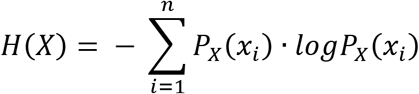

## RESULTS

### Chromatin occupancy profiling of replication origins throughout the cell cycle

We sought to profile the cell cycle-dependent changes in chromatin organization surrounding replication origins throughout the yeast genome. Cells were synchronized in late G1 using α-factor. Cells were then released from the α-factor arrest and samples were collected every 10 minutes for approximately two complete cell cycles (150 minutes) (Figure 1A). Biological replicates were performed, and the progression through the cell cycle was monitored by flow cytometry (Figure S1). Samples were aligned by cell cycle progression and the samples were merged for downstream analysis.

**Figure 1.**
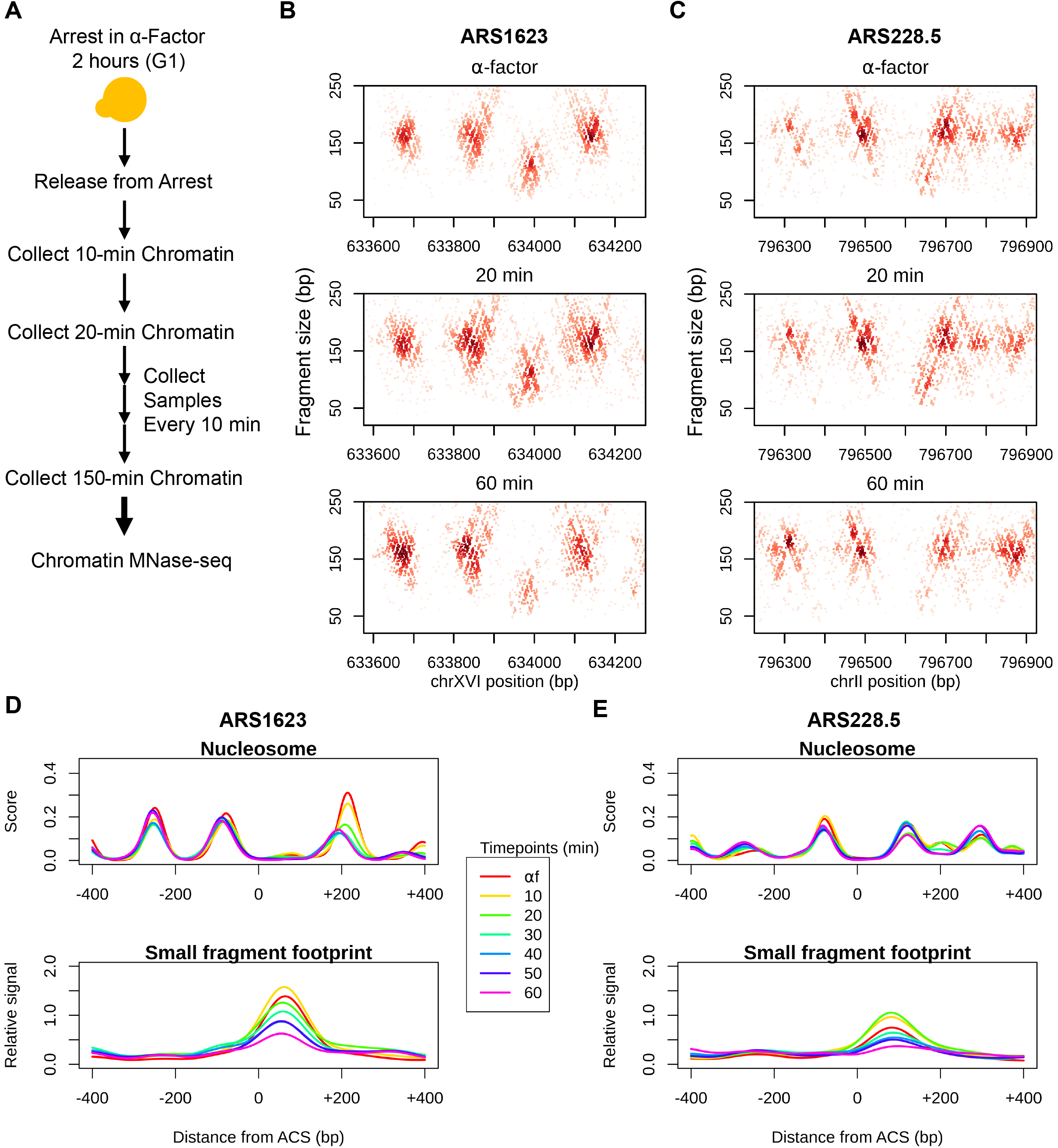
Profiling cell cycle-dependent chromatin dynamics by MNase mapping. (A) Schematic of the experimental design for capturing the cell cycle-dependent chromatin dynamics. (B) Chromatin profiles at *ARS1623* for select time points during the first cell cycle. The midpoints of recovered and sequenced MNase fragments are displayed. The size of each fragment is plotted as a function of its midpoint chromosomal position. (C) Chromatin profiles at *ARS228.5* for select time points during the first cell cycle. (D-E) Quantification of nucleosome scores and small fragment (< 120 bp) occupancy at all time points for the first cell cycle at *ARS1623* and *ARS228.5*, respectively. The same chromosome regions as in (B-C) are shown for each ARS locus.

To comprehensively interrogate chromatin dynamics at nucleotide resolution, we generated genome-wide chromatin occupancy profiles by digesting chromatin with MNase followed by paired-end sequencing [20,30,35]. DNA fragments protected by DNA binding factors (e.g., nucleosomes, transcription factors, ORC, etc.) are recovered and subjected to next-generation sequencing. The length and location of the mapped reads provide an unbiased view of chromatin occupancy throughout the genome. For example, nucleosomes protect DNA fragments of ~150 bp, and smaller DNA-binding factors (e.g., transcription factors and ORC) protect DNA fragments smaller than 120 bp. The mode of the distribution of fragment sizes recovered was 166 bp (Figure S2), consistent with the majority of DNA being packaged into nucleosomes throughout the genome. To visualize chromatin architecture at individual replication origins, we plotted the length of each fragment as a function of the chromosomal position of its midpoint. At *ARS1623*, an efficient and early origin, we observed an array of well-positioned nucleosomes flanking the origin, which were resolved as clusters of fragment midpoints centered at ~160 bp. We also observed an accumulation of smaller fragments at the ACS which represents an ORC-dependent footprint [20] (Figure 1B). Similar chromatin organization was observed at *ARS228.5*, an inefficient and late origin, albeit with “fuzzier” ACS-proximal nucleosomes, a narrower nucleosome free region (NFR), and a substantially weaker small fragment footprint at the ACS (Figure 1C).

We observed dynamic changes in chromatin organization as the cells proceeded through the cell cycle. As shown for select time points representing late G1 (α-factor), early S (20 min) and M (60 min) phases, we observed fluctuations in the dyad positions of the ACS-proximal nucleosomes and the occupancy of ACS-bound small fragments (Figures 1B and 1C). A score for nucleosome occupancy and position was calculated using a two-dimensional nucleosome kernel modeled on the MNase fragments associated with nucleosomes mapped by an orthogonal chemical cleavage method [32,33](Figure S3). A score for small factor occupancy (e.g. ORC, pre-RC and pre-IC) at origins was generated by calculating the density of small protected fragments less than 120 bp at the origin of DNA replication. To adjust for the variation in sample specific MNase digestion, we normalized the occupancy of the small fragment footprint at the ACS by the occupancy of the footprint at Abf1p binding sites (Figure S4). At the efficient *ARS1623*, the downstream (+1) nucleosome of the ACS was displaced further from the ACS at the α-factor time point compared with other time points in the first cell cycle (Figure 1D). The movement of the nucleosome away from the ACS in G1 is Cdc6-dependent and presumably facilitates pre-RC assembly [20]. In contrast, the +1 nucleosome of the inefficient *ARS228.5* appeared to be more static (Figure 1E). While the occupancy of ACS-bound small fragments is significantly stronger at *ARS1623*, both origins showed fluctuating small fragment occupancy throughout the cell cycle (Figures 1D and 1E), likely reflecting the cell cycle-coupled dynamics of helicase loading in G1 and the subsequent activation of the CMG holohelicase complex and movement away from the origin as the cell progresses into and through S-phase.

### Cell cycle-dependent changes in replication initiation factor occupancy at replication origins

The loading of the helicase to form the pre-RC and the recruitment of replication initiation factors at the origin are tightly coupled to the progression of the cell cycle. Each replication origin in the genome has an inherent efficiency which is thought to be determined, at least in part, by ORC binding, Mcm2-7 loading, the recruitment of activation factors, and the local chromatin environment [2,20,26,27,36,37]. We had previously used genome-wide chromatin occupancy profiling to identify ORC-dependent small fragment occupancy footprints at replication origins in two discrete phases of the cell cycle -- G1 and G2 [20]. In that study, we identified two classes of origins: the first class consisted of 264 origins which exhibited an ORC-dependent footprint in both G1 and G2 and a second, smaller (128) less efficient class of origins which exhibited an ORC-dependent footprint only in G1. To precisely quantify the dynamic changes in small fragment footprints at the ACS throughout the cell cycle, we calculated the small fragment density at the ACS of each origin with a bandwidth of 50 bp at each time point. We observed a periodic protection footprint at the ACS for both classes of origins (Figures 2A & B) which peaked in late G1/early S-phase in consecutive cell cycles and likely represents the recruitment of Cdc45 and GINS to the pre-RC to form the CMG holohelicase complex at the most efficient origins. After peaking in late G1/early S-phase, the density of small fragments at the ACS gradually declines to a nadir near mitosis. This decline in signal throughout S-phase likely reflects the activation and the bi-directional movement of the CMG holohelicase complex with the replication fork away from the origin and/or the disassembly of the pre-RC at any passively replicated origins. By late G2 and through mitosis it is likely that any remaining small fragments at the origins are due to ORC.

**Figure 2.**
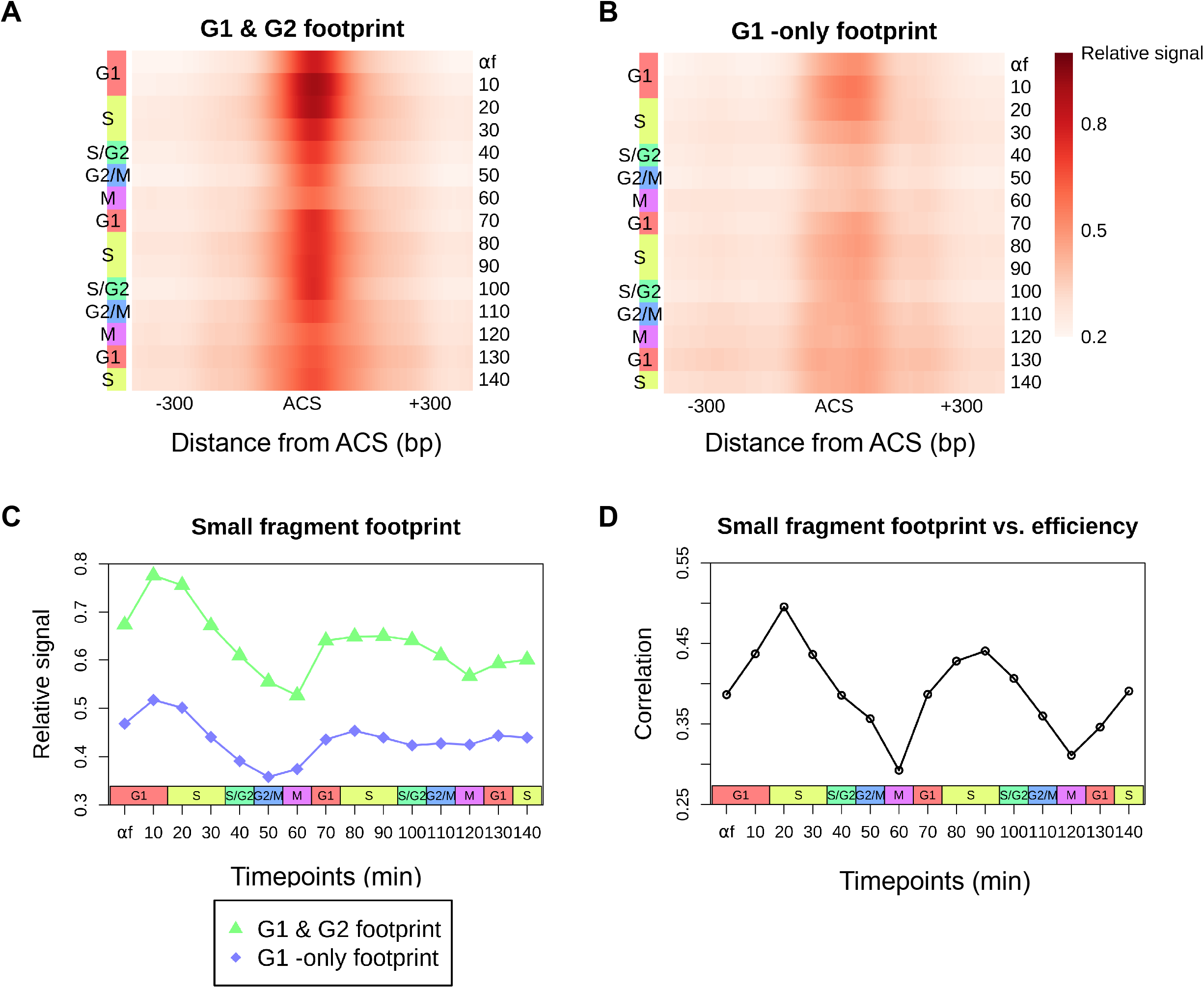
The cell cycle-dependent accumulation of small protected fragments at the ACS correlates with origin efficiency. (A-B) Heatmaps of aggregate small fragment (< 120 bp) occupancy at 264 origins with a previously described ORCdependent footprint in both “G1 and G2” (A) and 128 origins with an ORC-dependent footprint in “G1 only” (B) throughout the cell cycle [20]. All origins are oriented by the T-rich ACS strand. (C) Average small fragment occupancy +/− 100 bp surrounding the peak of the aggregate ORC-dependent footprint for each class of origins. (D) Spearman correlation between log2 transformed ACS-bound small fragment footprint density and the activation efficiency [2] for 371 origins exhibiting an ORC-dependent footprint at each time point.

The signal in the origins exhibiting an ORC-dependent footprint in “G1 only” was significantly dampened relative to the “G1&G2” class of origins (Figure 2C). Although we still observe an oscillatory pattern, the small fragment occupancy signals in G2 and M are barely at the detectable limit. Given the decreased signal observed throughout the cell cycle, this suggests a defect in either ORC recruitment or the stability of ORC on the DNA at these origins which ultimately leads to a stochastic defect in downstream helicase loading.

We previously found that stable ORC binding in both G1 and G2 was a determinant of efficient origins [20]. However, with our more holistic view of small fragment occupancy at the ACS throughout the cell cycle, we reasoned that perhaps a better predictor of origin efficiency might be the level of protected fragments at each origin as cells progress from G1 into S-phase. To test this hypothesis, we calculated the correlation between ACS-bound small fragment occupancy and origin efficiency at each time point throughout the cell cycle (Figure S5; Figure 2D). We found a cyclic pattern, with the correlation between origin efficiency and small fragment occupancy peaking in early S-phase and reaching its lowest point near mitosis. These results are consistent with recruitment of origin activation factors like Cdc45 and GINS to form the CMG holohelicase complex in late G1 and early S being a key determinant of origin activation.

### Cell cycle-regulated nucleosome occupancy dynamics around replication origins

The position of nucleosomes relative to the ACS impacts origin function [8,38–41]. We analysed the aggregate nucleosome signal of each origin class to profile the cell cycle-dependent dynamics of origin-flanking nucleosomes. For both classes, the conserved chromatin structure of replication origins - an NFR at the ACS surrounded by two well-positioned nucleosomes - was maintained throughout the cell cycle (Figure 3A).

**Figure 3.**
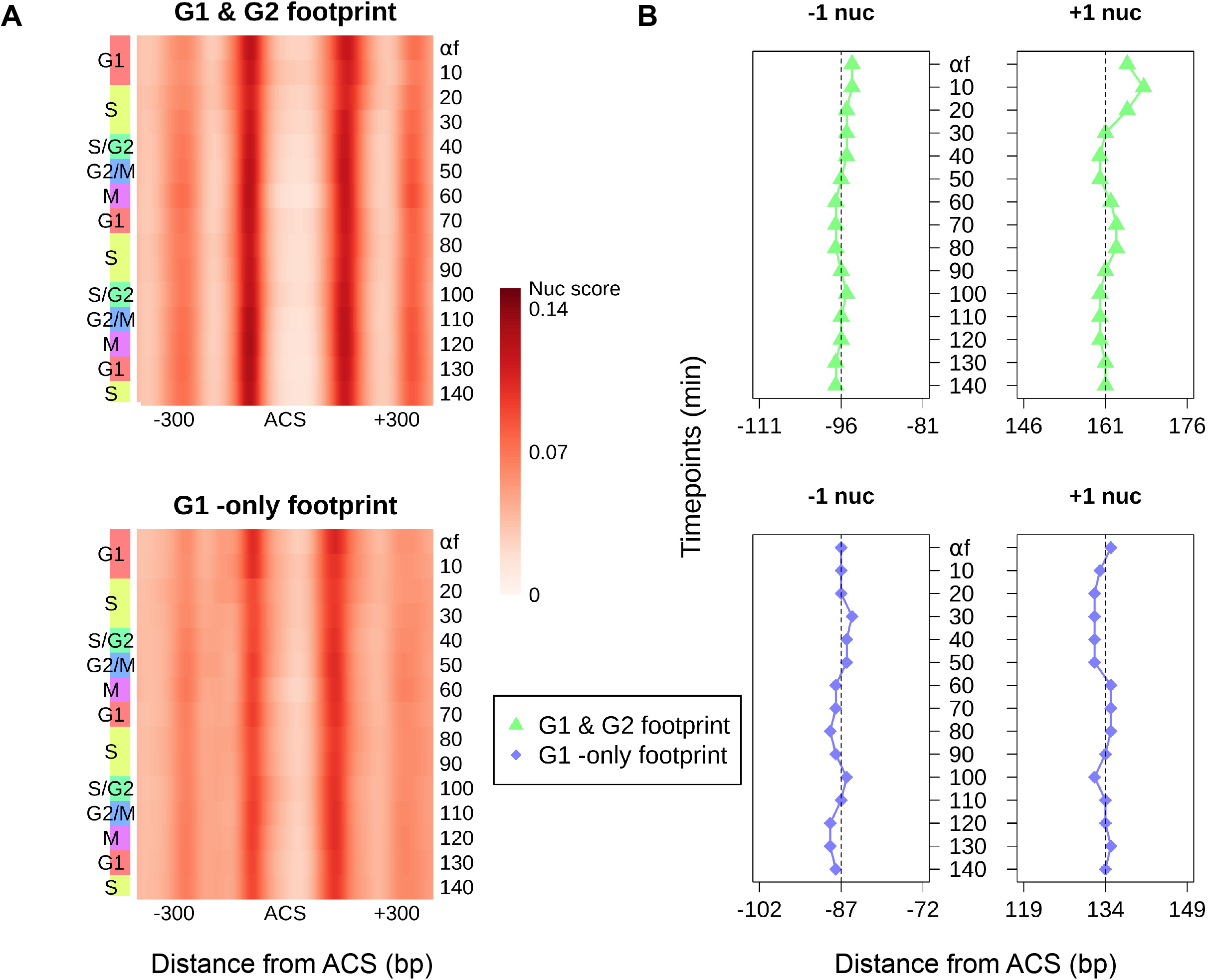
Cell cycle-dependent nucleosome dynamics at replication origins. (A) Heatmaps of aggregate nucleosome scores for 264 origins with an ORC-dependent footprint in both “G1 and G2” (top panel) and 128 origins with an ORC-dependent footprint in “G1 only” (bottom panel) through the cell cycle [20]. All origins are oriented by the T-rich ACS strand. (B) Dyad positions of the aggregate −1 and +1 nucleosomes relative to the ACS for each cell cycle time point and for each class of origins.

We first analysed the positioning of the NFR-proximal −1 and +1 nucleosomes throughout the cell cycle. We defined the dyad of either nucleosome as the position with maximal nucleosome score upstream or downstream of the ACS. The NFRs of origins with a footprint in both G1 and G2 are wider throughout the cell cycle (average dyad-to-dyad distance: 257.8 bp versus 220.9 bp). The increased width of the NFR was due to the downstream localization of the +1 nucleosome while the position of the −1 nucleosomes remained static (Figure 3B). Notably, the +1 nucleosome of origins with a G1 and G2 footprint exhibited periodic repositioning throughout the cell cycle, coincident with the helicase loading and recruitment of factors to form the Cdc45/Mcm2-7/GINS (CMG) complex. For example, the farthest +1 nucleosome positioning from the ACS was observed in the first G1/S transition (10 min), indicating recruitment of the CMG complex and pre-IC assembly. After origin activation, we observed that +1 nucleosome gradually started to move back towards the ACS with cell cycle progression, likely reflecting the subsequent dissociation of replication factors from origins. The extent of the nucleosome shift for the second cell cycle became weaker due to the gradual loss of cell synchrony. The +1 nucleosome positioning of origins with a “G1-only” footprint did not fluctuate in a cell cycle dependent manner which may be attributed to reduced formation of the rate-limiting CMG complex at less efficient origins [17,41].

### Replication-coupled nucleosome disorganization

The progression of the replication fork results in eviction of histones and disruption of nucleosome positioning [19]. We utilized Shannon entropy to assess the level of nucleosome organization [33] throughout the cell cycle. Well-positioned nucleosomes have a low entropy while disorganized nucleosomes exhibit a high entropy. We recently reported that helicase activation in the absence of primase activity resulted in the disorganization of origin proximal nucleosomes [29]. We reasoned that nucleosome disruption by helicase-induced unwinding would be a feature of sequences at the active replication fork. We calculated the entropy of nucleosome signals in 1kb windows for 30 kb surrounding the 69 most efficient origins (Figure 4A) and 69 most inefficient origins (Figure 4B). The mean entropy scores for each window were then ordered by their distance from the origin (rows) as a function of progression through the cell cycle (columns). In both the efficient and inefficient origins we observed increased entropy in S-phase consistent with disruption of chromatin by the passage of the replication fork. For the sequences most proximal to efficient origins, the entropy exhibited a temporal shift from early S (20 min) to mid S-phase (30 min), consistent with earlier timing of activation for efficient origins.

**Figure 4.**
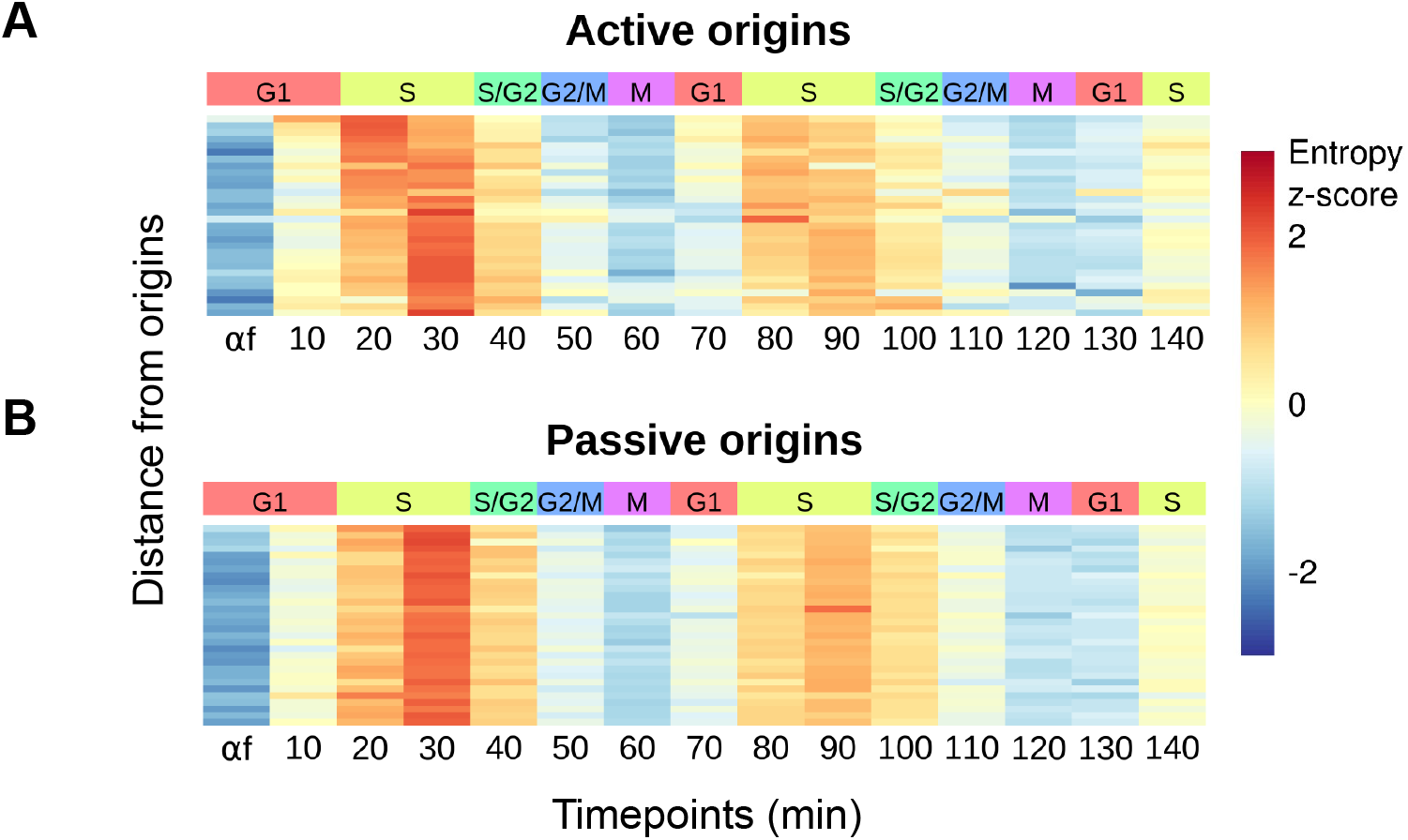
Nucleosome disruption at the replication fork. (A-B) Heatmaps representing the average nucleosome entropy in 1 kb windows for 30 kb surrounding the top 20% (n = 69) most efficient (active) origins (A) and the top 20% (n = 69) most inefficient (passive) origins (B) [2], among origins exhibiting an ORC-dependent footprint, at each time point. Each row represents a window and is ordered by the distance from the nearest origin. Nucleosome entropy is standardized into z-scores across each row.

## DISCUSSION

We interrogated chromatin dynamics at replication origins over the course of multiple consecutive cell cycles. As cells progressed through each cell cycle, we observed specific changes in chromatin occupancy that correspond to key steps in the selection and activation of start sites of DNA replication. We found that for each cell cycle, the downstream +1 nucleosome relative to the T-rich ACS moved further downstream in late G1 and early S-phase before returning back to its original position by G2. Similarly, we observed a cyclic increase in protected fragments spanning the ACS that likely represent pre-RC assembly and the recruitment of origin activation factors in late G1/early S. The accumulation of protected fragments at the ACS in early S-phase had the most predictive power for discerning active and efficient origins. Finally, we observed genome-wide disruption of nucleosomes traveling with active replication forks during S-phase, reflecting helicase induced DNA unwinding at the fork. Together, these data provide insights into the dynamic interplay between origin function and local chromatin environment with high spatiotemporal resolution.

In contrast to chromatin immunoprecipitation, genome-wide chromatin occupancy profiling by MNase digestion provides a holistic view of DNA protein occupancy that neither requires factor specific antibodies nor is encumbered by epitope accessibility [5]. A consequence of genome-wide chromatin occupancy being a factor agnostic assay is that it only reports if a DNA sequence is protected or occupied. The factor responsible for the occupancy has to be inferred from other sources such as sequence context (e.g. an ACS motif) or prior factor specific localization experiments. Thus, we need to infer that the subtle cell cycle-dependent changes in chromatin occupancy at the ACS reflect distinct steps in licensing and activation of replication origins. Importantly, in support of these assumptions, our prior work using static G1 and G2 arrested samples did demonstrate that the origin chromatin occupancy of the ACS was ORC dependent and that the increase in occupancy in G1 was dependent on origin licensing or pre-RC assembly [20].

The determinants of origin efficiency are poorly understood at the chromatin level. While it is clear that rate limiting activation factors establish the temporal order of origin activation [16,17], what is less clear is why and how individual origins are more or less sensitive to these factors. Local histone modifications have been shown to modulate replication timing with early work demonstrating that origin activity could be repressed by moving an active early origin to a heterochromatic region of the genome [42]. Global perturbation of histone acetylation levels also regulate genome-wide replication timing [10,43], but it is unclear if this is a direct or indirect effect via modulating the accessibility of the multicopy rDNA locus and sequestering origin activation factors [11]. The association of ORC with DNA can be driven by sequence or chromatin elements, with the chromatin-dependent ORC class being correlated with early replicating efficient origins [25]. The relative amount of Mcm2-7 loading in G1 is also predictive of origin function [27]. Our work identifies an increase in protected fragment occupancy at the ACS in early S-phase that is most predictive of efficient origin activation consistent with the formation of the CMG holohelicase complex at licensed origins. While this does not necessarily preclude earlier steps like ORC binding or Mcm2-7 loading as being deterministic, it is notable that the accumulation of protected fragments at the ACS in G2 (ORC alone) or G1 were less predictive of origin efficiency.

A consequence of the passage of the replication fork is disruption of chromatin ahead of the fork and the restoration of chromatin in the wake of the fork. Our chromatin occupancy profiling was able to capture the dynamic and transient disruption of chromatin associated with the replication fork. Specifically, we observed a transient and S-phase specific increase in entropy that was temporally linked to the distance from the nearest replication origin. The reassembly of nucleosomes and the chromatin landscape behind the fork is critical for epigenetic inheritance and factors that impair or delay assembly can significantly impact gene regulation [44–46] and differentiation [47]. Our ability to discern differences in entropy associated with replication coupled nucleosome assembly will enable future studies, with increased temporal resolution, to identify and characterize locus specific differences in chromatin assembly that may be governed in part by chromosome position, transcription and the local chromatin environment.

## Author Contributions

Conceptualization, Y.L., and D.M.; Formal analysis, Y.L; Funding acquisition, D.M. and A.H.; Project administration, D.M. and A.H.; Supervision, D.M. and A.H.; Writing original draft, Y.L.; Writing review & editing, A.H. and D.M. All authors have read and agreed to the published version of the manuscript.

## Data Availability Statement

All genomic data are publicly available at the NCBI GEO repository with the accession number GSE168699.

## ACKNOWLEDGMENTS

We acknowledge members of the MacAlpine Laboratory and the Hartemink Group for critical comments and suggestions during the development of this study. We thank Heather K. MacAlpine for critical reading and editing of this manuscript.

## REFERENCES

1. Lin, Y.C.; Prasanth, S.G. Replication initiation: Implications in genome integrity. DNA Repair (Amst) 2021, 103, 103131, doi:10.1016/j.dnarep.2021.103131.

2. McGuffee, S.R.; Smith, D.J.; Whitehouse, I. Quantitative, genome-wide analysis of eukaryotic replication initiation and termination. Mol Cell 2013, 50, 123–135, doi:10.1016/j.molcel.2013.03.004.

3. Aparicio, O.M. Location, location, location: it’s all in the timing for replication origins. Gene Dev 2013, 27, 117–128, doi:10.1101/gad.209999.112.

4. Wyrick, J.J.; Aparicio, J.G.; Chen, T.; Barnett, J.D.; Jennings, E.G.; Young, R.A.; Bell, S.P.; Aparicio, O.M. Genome-wide distribution of ORC and MCM proteins in S. cerevisiae: high-resolution mapping of replication origins. Science 2001, 294, 2357–2360, doi:10.1126/science.1066101.

5. Aparicio, O.M.; Weinstein, D.M.; Bell, S.P. Components and dynamics of DNA replication complexes in S. cerevisiae: redistribution of MCM proteins and Cdc45p during S phase. Cell 1997, 91, 59–69, doi:10.1016/s0092-8674(01)80009-x.

6. Retkute, R.; Nieduszynski, C.A.; de Moura, A. Dynamics of DNA replication in yeast. Phys Rev Lett 2011, 107, 068103, doi:10.1103/PhysRevLett.107.068103.

7. Muller, C.A.; Hawkins, M.; Retkute, R.; Malla, S.; Wilson, R.; Blythe, M.J.; Nakato, R.; Komata, M.; Shirahige, K.; de Moura, A.P.S.; et al. The dynamics of genome replication using deep sequencing. Nucleic Acids Res 2014, 42, doi:10.1093/nar/gkt878.

8. Eaton, M.L.; Galani, K.; Kang, S.; Bell, S.P.; MacAlpine, D.M. Conserved nucleosome positioning defines replication origins. Genes Dev 2010, 24, 748–753, doi:10.1101/gad.1913210.

9. Berbenetz, N.M.; Nislow, C.; Brown, G.W. Diversity of eukaryotic DNA replication origins revealed by genome-wide analysis of chromatin structure. Plos Genet 2010, 6, e1001092, doi:10.1371/journal.pgen.1001092.

10. Vogelauer, M.; Rubbi, L.; Lucas, I.; Brewer, B.J.; Grunstein, M. Histone acetylation regulates the time of replication origin firing. Mol Cell 2002, 10, 1223–1233, doi:10.1016/s1097-2765(02)00702-5.

11. Yoshida, K.; Bacal, J.; Desmarais, D.; Padioleau, I.; Tsaponina, O.; Chabes, A.; Pantesco, V.; Dubois, E.; Parrinello, H.; Skrzypczak, M.; et al. The histone deacetylases sir2 and rpd3 act on ribosomal DNA to control the replication program in budding yeast. Mol Cell 2014, 54, 691–697, doi:10.1016/j.molcel.2014.04.032.

12. Bell, S.P.; Stillman, B. ATP-dependent recognition of eukaryotic origins of DNA replication by a multiprotein complex. Nature 1992, 357, 128–134, doi:10.1038/357128a0.

13. Diffley, J.F.X.; Cocker, J.H.; Dowell, S.J.; Rowley, A. Two steps in the assembly of complexes at yeast replication origins in vivo. Cell 1994, 78, 303–316, doi:https://doi.org/10.1016/0092-8674(94)90299-2.

14. Bell, S.P.; Kaguni, J.M. Helicase Loading at Chromosomal Origins of Replication. Csh Perspect Biol 2013, 5, doi:10.1101/cshperspect.a010124.

15. Tanaka, S.; Araki, H. Helicase activation and establishment of replication forks at chromosomal origins of replication. Csh Perspect Biol 2013, 5, a010371–a010371, doi:10.1101/cshperspect.a010371.

16. Mantiero, D.; Mackenzie, A.; Donaldson, A.; Zegerman, P. Limiting replication initiation factors execute the temporal programme of origin firing in budding yeast. EMBO J 2011, 30, 4805–4814, doi:10.1038/emboj.2011.404.

17. Lynch, K.L.; Alvino, G.M.; Kwan, E.X.; Brewer, B.J.; Raghuraman, M.K. The effects of manipulating levels of replication initiation factors on origin firing efficiency in yeast. Plos Genet 2019, 15, e1008430, doi:10.1371/journal.pgen.1008430.

18. Xu, W.; Aparicio, J.G.; Aparicio, O.M.; Tavare, S. Genome-wide mapping of ORC and Mcm2p binding sites on tiling arrays and identification of essential ARS consensus sequences in S. cerevisiae. BMC Genomics 2006, 7, 276, doi:10.1186/1471-2164-7-276.

19. MacAlpine, D.M.; Almouzni, G. Chromatin and DNA replication. Cold Spring Harb Perspect Biol 2013, 5, a010207, doi:10.1101/cshperspect.a010207.

20. Belsky, J.A.; MacAlpine, H.K.; Lubelsky, Y.; Hartemink, A.J.; MacAlpine, D.M. Genome-wide chromatin footprinting reveals changes in replication origin architecture induced by pre-RC assembly. Genes Dev 2015, 29, 212–224, doi:10.1101/gad.247924.114.

21. Lubelsky, Y.; Sasaki, T.; Kuipers, M.A.; Lucas, I.; Le Beau, M.M.; Carignon, S.; Debatisse, M.; Prinz, J.A.; Dennis, J.H.; Gilbert, D.M. Pre-replication complex proteins assemble at regions of low nucleosome occupancy within the Chinese hamster dihydrofolate reductase initiation zone. Nucleic Acids Res 2011, 39, 3141–3155, doi:10.1093/nar/gkq1276.

22. Cayrou, C.; Coulombe, P.; Puy, A.; Rialle, S.; Kaplan, N.; Segal, E.; Mechali, M. New insights into replication origin characteristics in metazoans. Cell Cycle 2012, 11, 658–667, doi:10.4161/cc.11.4.19097.

23. Cayrou, C.; Ballester, B.; Peiffer, I.; Fenoui, R.; Coulombe, P.; Andrau, J.C.; van Helden, J.; Mechalil, M. The chromatin environment shapes DNA replication origin organization and defines origin classes. Genome Res 2015, 25, 1873–1885, doi:10.1101/gr.192799.115.

24. Raghuraman, M.K.; Winzeler, E.A.; Collingwood, D.; Hunt, S.; Wodicka, L.; Conway, A.; Lockhart, D.J.; Davis, R.W.; Brewer, B.J.; Fangman, W.L. Replication dynamics of the yeast genome. Science 2001, 294, 115–121, doi:10.1126/science.294.5540.115.

25. Hoggard, T.; Shor, E.; Muller, C.A.; Nieduszynski, C.A.; Fox, C.A. A Link between ORC-Origin Binding Mechanisms and Origin Activation Time Revealed in Budding Yeast. Plos Genet 2013, 9, doi:10.1371/journal.pgen.1003798.

26. Das, S.P.; Borrman, T.; Liu, V.W.T.; Yang, S.C.H.; Bechhoefer, J.; Rhind, N. Replication timing is regulated by the number of MCMs loaded at origins. Genome Res 2015, 25, 1886–1892, doi:10.1101/gr.195305.115.

27. Dukaj, L.; Rhind, N. The capacity of origins to load mcm establishes replication timing patterns. Plos Genet 2021, 17, doi:10.1371/journal.pgen.1009467.

28. Speck, C.; Chen, Z.; Li, H.; Stillman, B. ATPase-dependent cooperative binding of ORC and Cdc6 to origin DNA. Nat Struct Mol Biol 2005, 12, 965–971, doi:10.1038/nsmb1002.

29. Hoffman, R.A.; MacAlpine, H.K.; MacAlpine, D.M. Disruption of origin chromatin structure by helicase activation in the absence of DNA replication. Genes Dev 2021, 35, 1339–1355, doi:10.1101/gad.348517.121.

30. Henikoff, J.G.; Belsky, J.A.; Krassovsky, K.; MacAlpine, D.M.; Henikoff, S. Epigenome characterization at single base-pair resolution. Proc Natl Acad Sci U S A 2011, 108, 18318–18323, doi:10.1073/pnas.1110731108.

31. Langmead, B.; Trapnell, C.; Pop, M.; Salzberg, S.L. Ultrafast and memory-efficient alignment of short DNA sequences to the human genome. Genome Biology 2009, 10, doi:10.1186/gb-2009-10-3-r25.

32. Brogaard, K.R.; Xi, L.Q.; Wang, J.P.; Widom, J. A Chemical Approach to Mapping Nucleosomes at Base Pair Resolution in Yeast. Method Enzymol 2012, 513, 315–334, doi:10.1016/B978-0-12-391938-0.00014-8.

33. Tran, T.Q.; MacAlpine, H.K.; Tripuraneni, V.; Mitra, S.; MacAlpine, D.M.; Hartemink, A.J. Linking the dynamics of chromatin occupancy and transcription with predictive models. Genome Res 2021, 31, 1035–1046, doi:10.1101/gr.267237.120.

34. MacIsaac, K.D.; Wang, T.; Gordon, D.B.; Gifford, D.K.; Stormo, G.D.; Fraenkel, E. An improved map of conserved regulatory sites for Saccharomyces cerevisiae. BMC Bioinformatics 2006, 7, 113, doi:10.1186/1471-2105-7-113.

35. Kent, N.A.; Adams, S.; Moorhouse, A.; Paszkiewicz, K. Chromatin particle spectrum analysis: a method for comparative chromatin structure analysis using paired-end mode next-generation DNA sequencing. Nucleic Acids Res 2011, 39, e26, doi:10.1093/nar/gkq1183.

36. Hawkins, M.; Retkute, R.; Muller, C.A.; Saner, N.; Tanaka, T.U.; de Moura, A.P.S.; Nieduszynski, C.A. High-Resolution Replication Profiles Define the Stochastic Nature of Genome Replication Initiation and Termination. Cell Reports 2013, 5, 1132–1141, doi:10.1016/j.celrep.2013.10.014.

37. Muller, P.; Park, S.; Shor, E.; Huebert, D.J.; Warren, C.L.; Ansari, A.Z.; Weinreich, M.; Eaton, M.L.; MacAlpine, D.M.; Fox, C.A. The conserved bromo-adjacent homology domain of yeast Orc1 functions in the selection of DNA replication origins within chromatin. Gene Dev 2010, 24, 1418–1433, doi:10.1101/gad.1906410.

38. Simpson, R.T. Nucleosome Positioning Can Affect the Function of a Cis-Acting DNA Element Invivo. Nature 1990, 343, 387–389, doi:DOI 10.1038/343387a0.

39. Lipford, J.R.; Bell, S.P. Nucleosomes positioned by ORC facilitate the initiation of DNA replication. Mol Cell 2001, 7, 21–30, doi:Doi 10.1016/S1097-2765(01)00151-4.

40. Berbenetz, N.M.; Nislow, C.; Brown, G.W. Diversity of Eukaryotic DNA Replication Origins Revealed by Genome-Wide Analysis of Chromatin Structure. Plos Genet 2010, 6, doi:10.1371/journal.pgen.1001092.

41. Azmi, I.F.; Watanabe, S.; Maloney, M.F.; Kang, S.; Belsky, J.A.; MacAlpine, D.M.; Peterson, C.L.; Bell, S.P. Nucleosomes influence multiple steps during replication initiation. Elife 2017, 6, doi:10.7554/eLife.22512.

42. Ferguson, B.M.; Fangman, W.L. A position effect on the time of replication origin activation in yeast. Cell 1992, 68, 333–339, doi:10.1016/0092-8674(92)90474-q.

43. Knott, S.R.; Viggiani, C.J.; Tavare, S.; Aparicio, O.M. Genome-wide replication profiles indicate an expansive role for Rpd3L in regulating replication initiation timing or efficiency, and reveal genomic loci of Rpd3 function in Saccharomyces cerevisiae. Genes Dev 2009, 23, 1077–1090, doi:10.1101/gad.1784309.

44. Huang, S.B.; Zhou, H.; Katzmann, D.; Hochstrasser, M.; Atanasova, E.; Zhang, Z.G. Rtt106p is a histone chaperone involved in heterochromatin-mediated silencing. P Natl Acad Sci USA 2005, 102, 13410–13415, doi:10.1073/pnas.0506176102.

45. Kaufman, P.D.; Kobayashi, R.; Stillman, B. Ultraviolet radiation sensitivity and reduction of telomeric silencing in Saccharomyces cerevisiae cells lacking chromatin assembly factor-I. Genes Dev 1997, 11, 345–357, doi:10.1101/gad.11.3.345.

46. Meijsing, S.H.; Ehrenhofer-Murray, A.E. The silencing complex SAS-I links histone acetylation to the assembly of repressed chromatin by CAF-I and Asf1 in Saccharomyces cerevisiae. Genes Dev 2001, 15, 3169–3182, doi:10.1101/gad.929001.

47. Escobar, T.M.; Oksuz, O.; Saldana-Meyer, R.; Descostes, N.; Bonasio, R.; Reinberg, D. Active and Repressed Chromatin Domains Exhibit Distinct Nucleosome Segregation during DNA Replication. Cell 2019, 179, 953–963 e911, doi:10.1016/j.cell.2019.10.009.

