## Supplemental Materials for "Cell cycle-dependent chromatin dynamics at replication origins"

### SUPPLEMENTAL FIGURE LEGENDS

**Figure S1. Flow cytometry profiles monitoring cell cycle progression for each biological duplicate.** The distribution of DNA content for 30,000 cells is plotted for each sample.

**Figure S2. Distribution of MNase fragment lengths after subsampling and merging replicates.** Fragments shorter than 120 bp are considered small fragments and are notated by the red dashed line. The blue dashed line represents the mode of fragment sizes (166 bp).

**Figure S3. Density plot of the two-dimensional kernel modeled from an orthogonal set of nucleosome dyads.** The kernel is constructed from the aggregate MNase signal of 8,632 unique nucleosome positions on ChrIV previously mapped by a sensitive chemical cleavage method [32]. The kernel of the α-factor time point is shown.

**Figure S4. Small fragment footprint occupancy at Abf1p binding sites throughout the cell cycle.** (A) Heatmap showing unnormalized aggregate small fragment footprint density for 151 Abf1p binding sites at each time point. Subtle fluctuations in Abf1p footprint signal at different time points (but not correlated with cell cycle) are evident, likely reflecting sample specific variations in MNase digestion. (B) Heatmap representing Abf1p-normalized small fragment occupancy as in (A).

**Figure S5. Correlation between small factor occupancy at the ACS and origin efficiency.** Scatter plots of the activation efficiency [2] and log2 transformed ACS-bound small fragment density for individual origins exhibiting an ORC-dependent footprint (n = 371) at each time point. Spearman correlation is calculated for each time point.

Li et al., Figure S1

Replicate 1

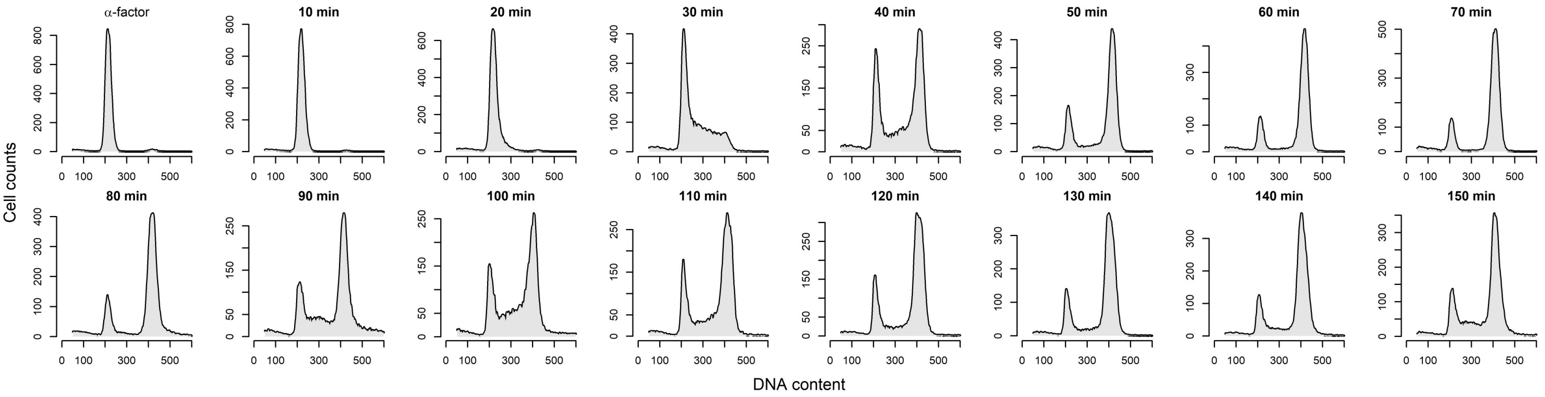


Replicate2

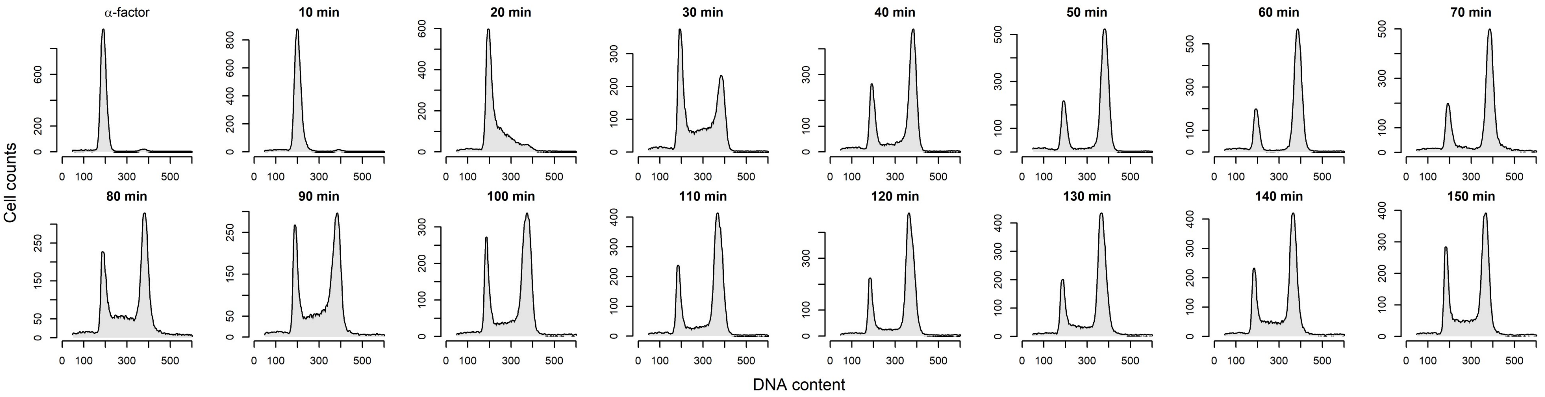

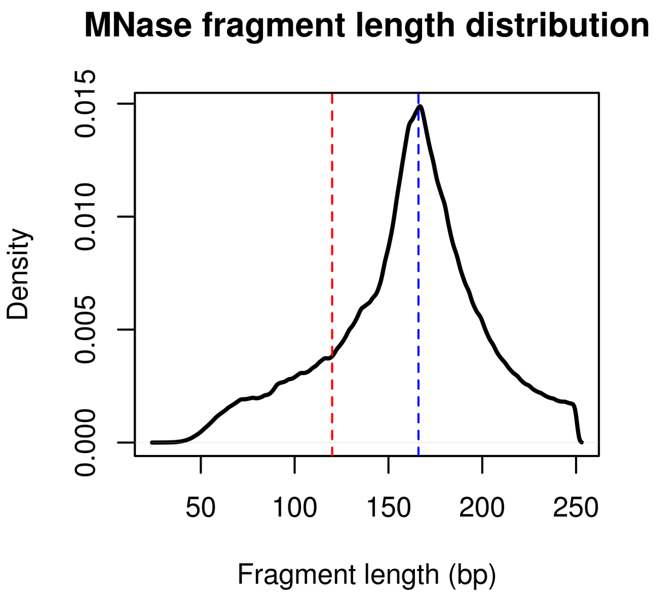
MNase fragment length distribution
Density
0.000
0.005
0.010
0.015
50
100
150
200
250
Fragment length (bp)

Li et al., Figure S3

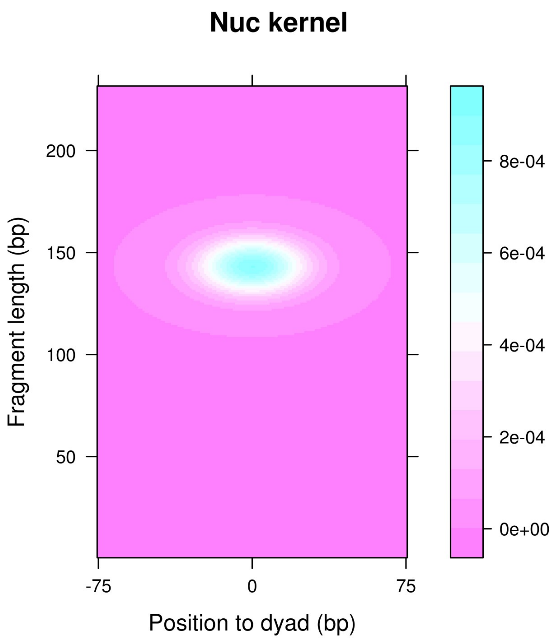

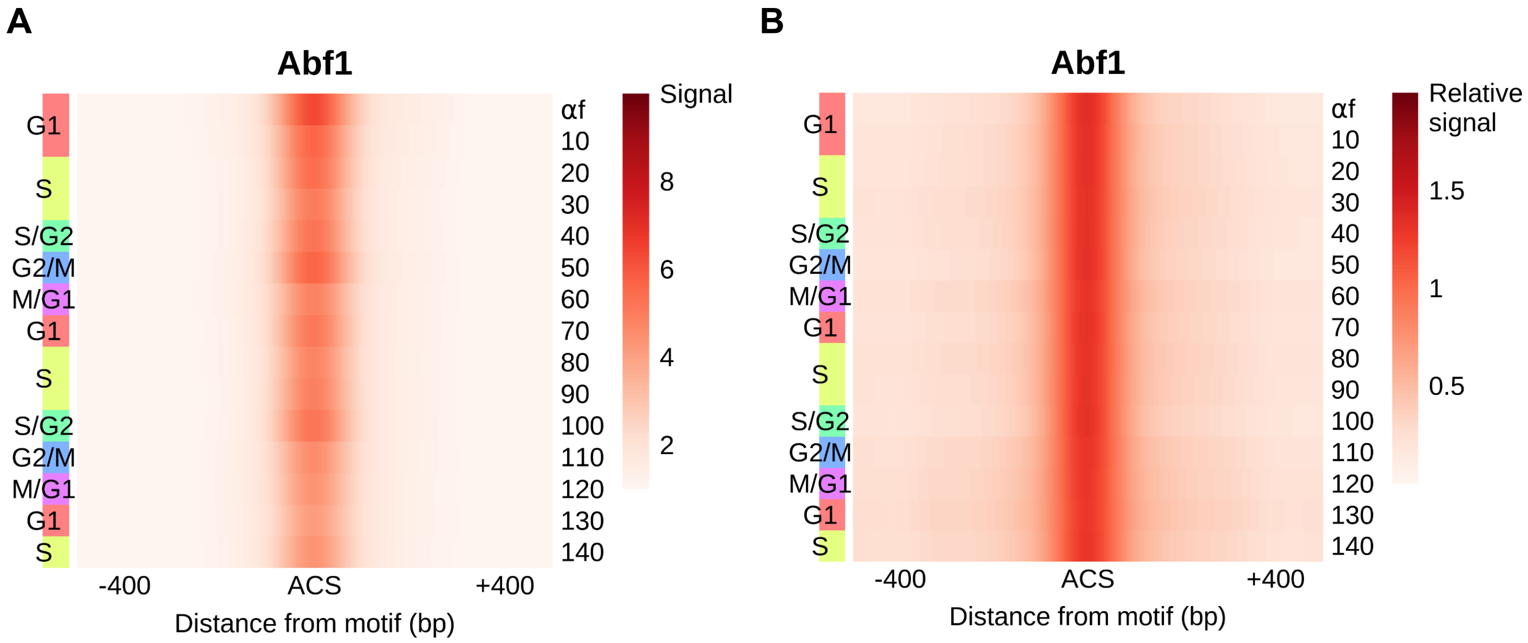
A
Abf1
G1
S
S/G2
G2/M
M/G1
G1
S
S/G2
G2/M
M/G1
G1
S
αf
10
20
30
40
50
60
70
80
90
100
110
120
130
140
Signal
8
6
4
2
-400
ACS
+400
Distance from motif (bp)
B
Abf1
G1
S
S/G2
G2/M
M/G1
G1
S
S/G2
G2/M
M/G1
G1
S
αf
10
20
30
40
50
60
70
80
90
100
110
120
130
140
Relative
signal
1.5
1
0.5
-400
ACS
+400
Distance from motif (bp)

Li et al., Figure S5

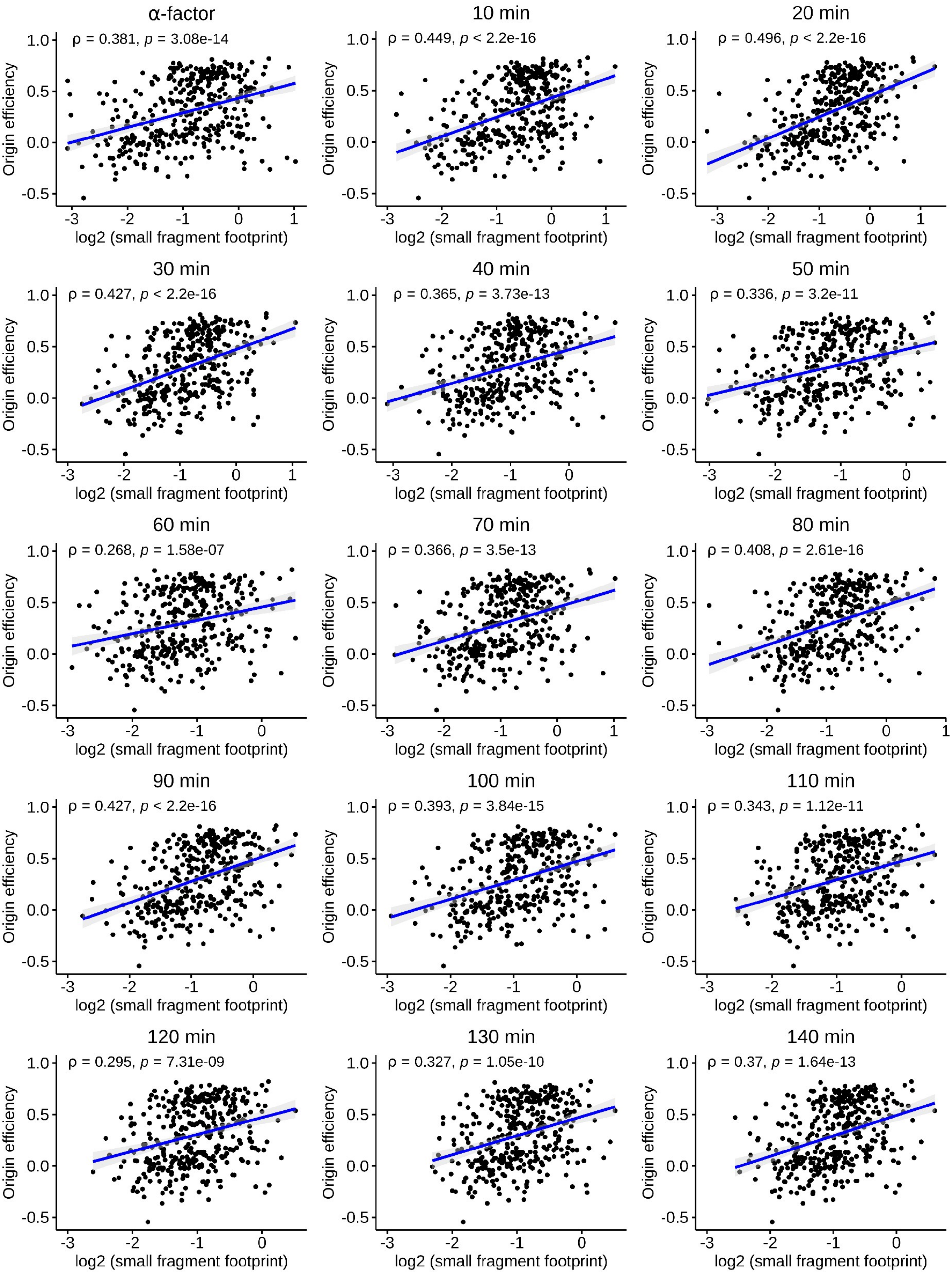
